# Full-Length Single-Molecule Protein Fingerprinting

**DOI:** 10.1101/2023.09.26.559471

**Authors:** Mike Filius, Raman van Wee, Carlos de Lannoy, Ilja Westerlaken, Zeshi Li, Sung Hyun Kim, Cecilia de Agrela Pinto, Yunfei Wu, Geert-Jan Boons, Martin Pabst, Dick de Ridder, Chirlmin Joo

## Abstract

Proteins are the primary functional actors of the cell. Hence, their identification is pivotal to advance our understanding of cell biology and disease. Current protein analysis methods are of limited use for distinguishing proteoforms. In particular, mass spectrometric methods often provide only ambiguous information on post-translational modification sites, and sequences of co-existing modifications may not be resolved. Here we demonstrate FRET-based single-molecule protein fingerprinting to map the location of individual amino acids and a post-translational modification within single full-length protein molecules. Using an approach that relies on transient binding of fluorescently labeled DNA strands to probe the amino acids on a protein one by one we show that we can fingerprint intrinsically disordered proteins as well as folded globular proteins with sub-nanometer resolution. We anticipate that this technology will be used for proteoform identification in biological and translational research with ultimate sensitivity.

## Introduction

Protein synthesis is a highly complex and regulated process, and much of this regulation occurs beyond the genome and transcriptome level. Via mechanisms such as alternative splicing and post-translational modifications (PTMs), a single protein encoding gene can produce hundreds of unique protein products, or proteoforms.^1^ Even subtle differences between proteoforms can dramatically alter their biological functioning and the expression of aberrant proteoforms is implicated in many diseases, including neurodegenerative diseases, metabolic disorders and a variety of cancers.^2–4^ It is increasingly appreciated that protein functionalities in a given biological context need to be analyzed at proteoform level, rather than at the coding gene level.

The proteoform information can only be obtained without fault when the protein of interest is studied intact. For example, specific proteoforms arising from phosphorylation or alternative splicing are conventionally detected with affinity-based approaches using probes (e.g. antibodies).^5,6^ However, these approaches may suffer from low specificity and are limited by the number of probes that are available. Recently, high-resolution native mass spectrometry (MS) has shown to be a powerful approach to investigate proteoform profiles.^7,8^ However, exact information on the sequence of co-occurring modifications cannot be determined by the widely employed bottom-up approaches. Alternative top-down fragmentation experiments require large sample quantities, purity and significant data interpretation, and may not be applicable to isobaric proteoforms without additional separation efforts.^9^ Analyzing full-length proteins at single-molecule resolution will offer a powerful solution to issues with the existing approaches.

While single-molecule sequencing of DNA and RNA is omnipresent^10,11^, the nature of proteins creates several challenges that have thus far precluded their sequencing at the single-molecule level.^12–15^ The increased number of building blocks in the polymer backbone from 4 nucleobases to 20 different amino acids complicates their discrimination and hinders specific labeling. The protein sequencing task is further impeded by the absence of a polymerase-like enzyme that can replicate proteins. Thirdly, protein folding and interactions are much less predictable than nucleic acid basepairing. As a workaround for these challenges, multiple groups have proposed protein fingerprinting, in which partial sequence information is used to generate a unique protein fingerprint.^16–20^ By mapping this fingerprint against a reference database, a protein can be identified. Thus far, proof-of-concept studies for protein fingerprinting have been limited to small model peptides,^19,21–23^ as their feasibility for full-length protein is often hampered by their resolution, throughput or experimental complexity.

Here we use our single-molecule protein fingerprinting technology, termed FRET by DNA eXchange, or FRET X, in which the distances of multiple specific amino acid residues to a reference point on an intact protein are measured via FRET (Förster Resonance Energy Transfer).^19,24^ These nanoscale distances are inferred from the FRET efficiency and constitute the unique protein fingerprint, allowing for the identification of the protein analytes. Central to this technology is the use of fluorescently labeled short DNA oligos that transiently bind to the complementary sequence conjugated to specific amino acid residues of the protein. The use of short DNA strands for protein fingerprinting has four main advantages: (1) the transient binding of the DNA probes allows for the detection of a single FRET pair at a time, even when multiple points of interest are present, which is not possible by directly labeling the amino acids of interest with fluorophores; (2) the highly specific and programmable nature of DNA hybridization allows for the specific and controlled targeting of each target residue (e.g. amino acid or PTM), much alike the super-resolution imaging technique DNA-PAINT^25,26^; (3) the pool of fluorescently labeled DNA probes are constantly replenished, eliminating concerns over photobleaching and enabling indefinite signal collection; and (4) the repeated interrogation of the same FRET pairs on a protein increases the fingerprinting precision.

We demonstrate that full-length and folded proteins can be analysed with FRET X and highlight its high localization precision. Harnessing the high resolving power, we demonstrate the ability of our full-length single-molecule protein fingerprinting technology to map PTM sites in intact proteins. We further show that with site-specific N-terminal bioconjugation, FRET X can be extended to non-recombinantly tagged proteins, a critical step towards analyzing native samples.

## Results

### Single-molecule fingerprinting

To demonstrate the concept of protein fingerprinting using FRET X, we designed a single-molecule FRET assay where a DNA labeled protein is immobilized on a PEGylated surface in a microfluidic device through biotin-streptavidin conjugation (**Fig. 1a**). This single-stranded DNA (ssDNA) at the protein terminus is used to immobilize the protein and also functions as a docking site for transient binding of complementary acceptor (Cy5)-labeled imager strands. The replenishment of both the donor and acceptor fluorophores through transient binding of their respective imager strands allows for extended periods of imaging, thereby enabling us to collect sufficient FRET events to determine the protein fingerprint with high precision.^24^ The cysteine residues introduced at different positions were labeled with an orthogonal DNA sequence to allow transient binding of donor (Cy3)-labeled imager strands (**Fig. 1a and Supplementary Fig. 1a**). The donor and acceptor imager strands were designed to have mean dwell times (Δτ) of 0.5 ± 0.1 s and 2.1 ± 0.1 s (**Supplementary Fig. 1b**,**c**), respectively. Binding of both imager strands was sufficiently weak to ensure dissociation and thereby repetitive, transient binding, but it was long enough to allow precise determination of the FRET efficiency for several minutes (**Supplementary Fig. 1d**).^24^ Furthermore, to increase the probability of the presence of the acceptor imager strand upon donor imager strand binding and thus allow for FRET, we injected 5-fold molar excess of the acceptor imager strand over the donor imager strand.

**Fig. 1:**
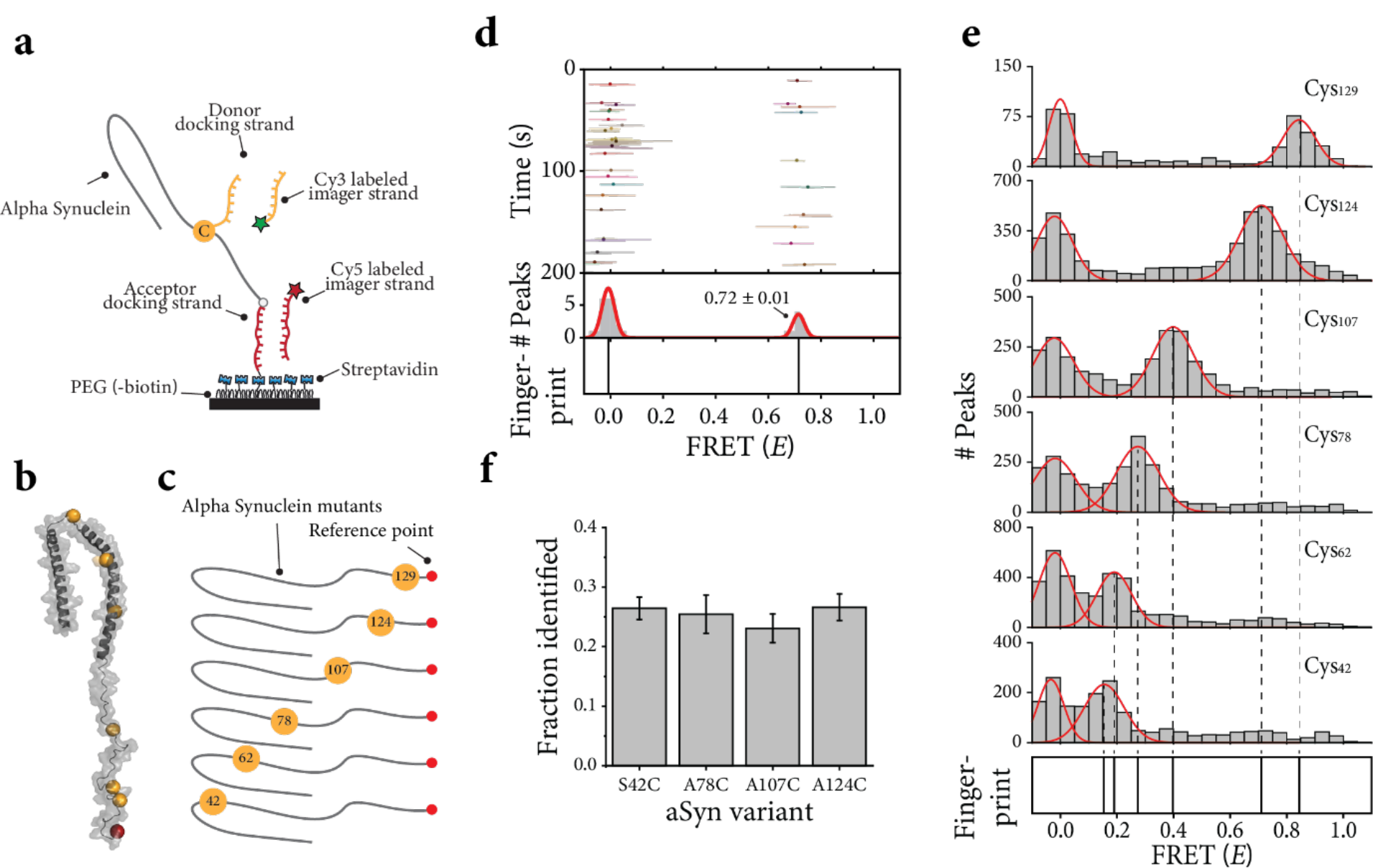
Repetitive binding of short DNA imager strands allows for high-resolution protein fingerprinting. (**a**) Schematic representation of the single-molecule assay. The model protein, alpha-synuclein (aSyn), is conjugated to a biotinylated single-stranded DNA (ssDNA) strand (red) to facilitate immobilization of the target protein to the PEGylated quartz surface. The donor (Cy3) labeled imager strand (yellow) binds to the DNA docking site on the cysteine, while the acceptor (Cy5) labeled imager strand (red) hybridizes to the docking site at the C-terminus of the protein. Simultaneous binding generates short FRET events and these are observed with total internal reflection microscopy. (**b**) Three-dimensional conformation for micelle-bound alpha-synuclein (PDB entry 1XQ8) with the location of the six cysteines probed (orange) and the C-terminus (red) indicated. (**c**) Schematic representation of the six aSyn constructs. Each construct contains a single cysteine (orange circle), whose position relative to the N-terminus is indicated, and a reference point at the C-terminus (red circle). (**d**) Representative kymograph from a single aSyn protein with a cysteine (Cys_124_) 18 amino acids separated from the reference point. The FRET efficiency for each data point in a binding event (lines) and the mean FRET efficiency from all data points in a binding event (dots) are indicated over the course of an experiment. The distribution of the average FRET efficiencies per FRET event is fitted with a Gaussian function. The mean values of the Gaussian fits are plotted in a separate panel (bottom) and are referred to as the FRET X fingerprint of the protein. The population on the left (*E ∼* 0) originates from events where the acceptor fluorophore was absent. The mean ± SEM of the Gaussian fit of the FRET peak are indicated in the plot. (**e**) Ensemble FRET X histograms for each of the aSyn constructs shown in panel c (single-molecule and ensemble kymographs shown in **Supplementary Fig. 2**), the mean ± FWHM FRET efficiencies were 0.84 ± 0.13 for Cys_129_, 0.71 ± 0.18 for Cys_124_, 0.40 ± 0.17 for Cys_107_, 0.27 ± 0.17 for Cys_78_, 0.19 ± 0.14 for Cys_62_, 0.15 ± 0.12 for Cys_42_. (**f**) Relative frequencies of detection for equimolar mixtures of four aSyn contstructs, as determined by the trained SVM. The error bars represent the 95% confidence interval over 1000 bootstrap iterations.

To demonstrate the ability of FRET X to fingerprint proteins, we constructed six human alpha-synuclein (aSyn) model proteins (**Fig. 1b,c**). Each variant contains a genetically introduced cysteine and has a biotinylated ssDNA strand conjugated to its C-terminus via an aldehyde encoding sequence.^27^ The different aSyn proteins were designed to have a varying distance between their cysteine and the reference point. We constructed a kymograph with the FRET events for each single protein molecule (**Fig. 1d and Supplementary Fig. 2 and 3**), where the lines indicate the FRET efficiency (*E*) for each data point and the dots are the mean FRET efficiency per event. The mean FRET efficiencies were fitted with a Gaussian mixture model (GMM) and we used the Bayesian information criterion to select the best number of distributions to fit each histogram. The Gaussian function was used to resolve the center of each peak with high precision (**Supplementary Fig. 1e,f**). The center values of each peak in the histogram are plotted in a separate panel, and this panel constitutes the protein fingerprint (**Fig. 1d,e**, bottom panel). The precision of the fingerprint is dependent on the number of binding events, and we can experimentally determine the fingerprint with a precision of Δ*E* ∼ 0.03 after 10 binding events (**Supplementary Fig. 1g**), underscoring the benefit of our DNA hybridization scheme in which the impact of stochastic photophysical effects is mitigated through repeated probing. It should be noted that the standard integration time of our measurement is 100 ms, which is several orders of magnitude slower than the typical time scale of protein conformational dynamics^28^, and thus we expect only a single FRET peak per point of interest.

For each aSyn variant we observed repetitive binding of the imager strand at the single-molecule level (**Supplementary Fig. 2 & 3**), and this yielded clearly distinct distributions and fingerprints, with the FRET efficiency monotonously decreasing as the distance to the C-terminal reference point increases (**Fig. 1e**). This experiment shows that FRET X has a range of ∼100 amino acids and that target amino acids whose location differ by 5 amino acids (Cys_124_ and Cys_129_) are still discernible. We next sought to determine the classification accuracy of different aSyn constructs, all added in equal proportions to a mixture. To accomplish this, a support vector machine (SVM) classifier was first trained on FRET values obtained from four separate experiments, each containing a single aSyn mutant. The trained SVM was then used to classify individual molecules within the mixture on the basis of their respective FRET values, enabling us to determine the relative concentrations of each of the constructs (**Supplementary Fig. 4**). We demonstrate that we are able to retrieve the initial relative abundance of each construct with high reproducibility (**Fig. 1f**).

### Single-molecule fingerprinting of disordered proteins

As a single type of amino acid can recur multiple times in a protein sequence, FRET X fingerprinting requires the detection of multiple FRET pairs in a single protein. To demonstrate that the transient and repetitive nature of binding events in FRET X facilitates fingerprinting of species with multiple FRET pairs, we designed two aSyn constructs, each containing two cysteines. The distance between the reference point and the first cysteine (Cys_124_) is identical for both constructs, while the distance to the second cysteine differs by 21 amino acids (Cys_78_ for **Fig. 2a**, and Cys_99_ for **Fig. 2b**). Upon performing our FRET X fingerprinting assay, we observed two distinct FRET populations for both constructs (**Fig. 2a,b and Supplementary Fig. 5a,b**). We observed a high FRET peak reporting on the relative position of Cys_124_, that was similar for both constructs (**Fig. 2a,b**). As expected, the average FRET efficiency of the second cysteine differs between Cys_78_ (0.32, **Fig. 2a**) and Cys_99_ (0.43, **Fig. 2b**). Furthermore, the FRET efficiencies for the double cysteine constructs are similar to the FRET efficiencies found in our experiments with the single cysteine constructs, demonstrating the reproducibility of FRET X protein fingerprinting (**Fig. 1e,f**).

**Fig. 2:**
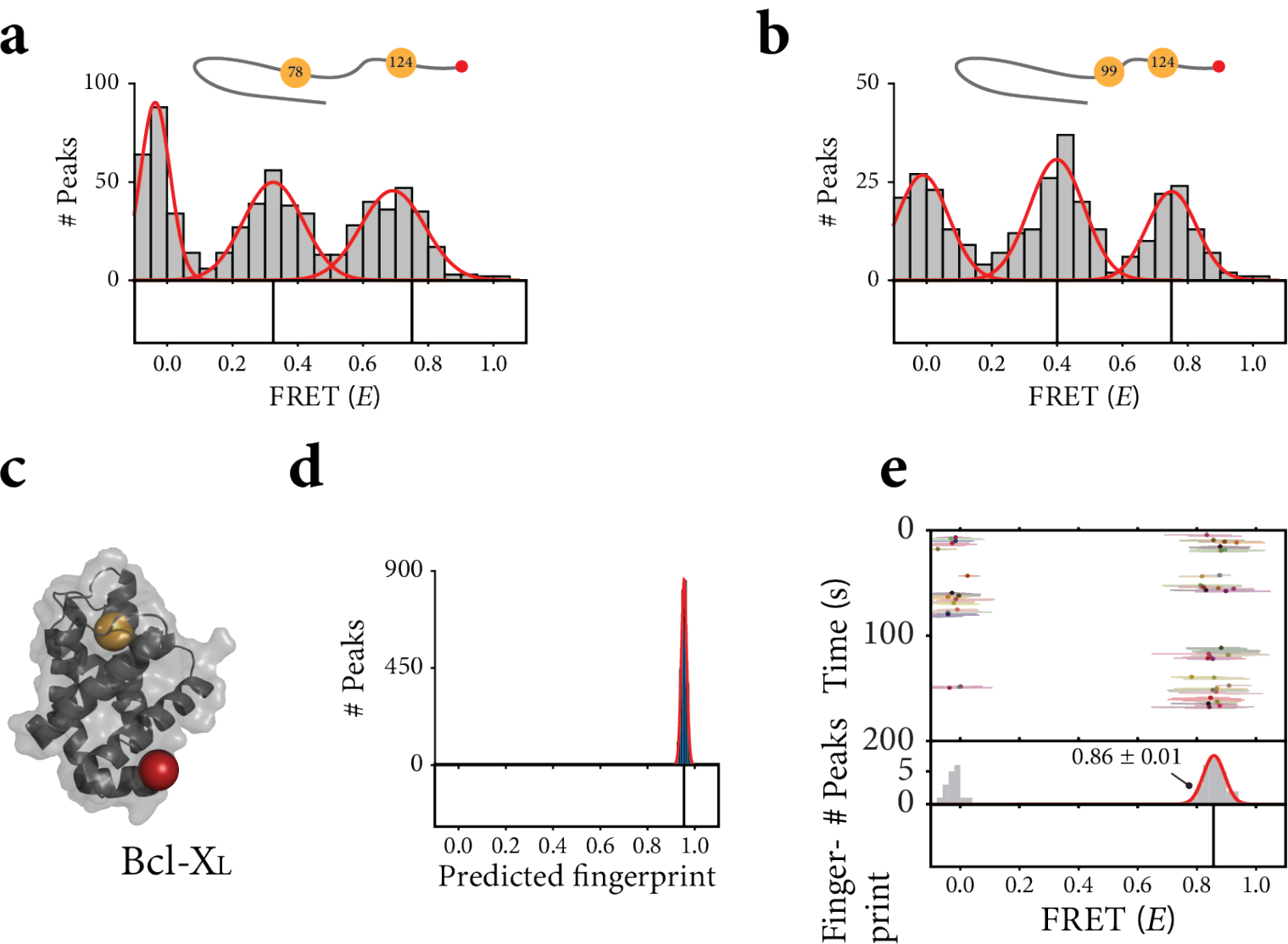
Single-molecule protein fingerprinting of disordered and folded proteins. (**a-b**) Top panels are schematic representations of the double cysteine variants of the aSyn model substrate. The cysteines (orange circles) are labeled with a DNA donor docking strand, and the C-terminus (red circle) is labeled with a DNA acceptor docking strand. All constructs contain a cysteine at position 124, with the second cysteine at varying positions. Middle panels; ensemble distributions of the observed FRET events. Bottom panels; FRET X histograms and fingerprints reporting on the relative distance of the cysteines to the reference point. The mean ± FWHM of the Gaussian fits for aSyn-Cys_78_+Cys_124_ (**a**) are 0.32 ± 0.10 and 0.70 ± 0.18 and for aSyn-Cys_99_+Cys_124_ (**b**) 0.43 ± 0.14 and 0.77 ± 0.16. (**c**) Three-dimensional conformation for BCLX_L_ (PDB entry: 1R2D) with the cysteine indicated in orange. (**d)** Predicted fingerprint (mean ± FWHM) for Bcl-X_L_ (**f**, 0.95 ± 0.03).The predicted fingerprint histograms were built from simulated FRET efficiencies from 200 individual molecules each having 10 FRET events. (**e)** Representative single-molecule kymograph with its determined fingerprint (mean ± SEM) of 0.86 ± 0.01 for Bcl-X_L_.

### Single-molecule fingerprinting of globular proteins

To effectively fingerprint cellular proteins at the single-molecule level, our FRET X platform should be able to cope with the folded structure that most cellular proteins have. To demonstrate FRET X for single-molecule fingerprinting of folded proteins, we purified the human apoptosis regulator Bcl-2-like protein 1 (Bcl) isoform Bcl-X_L_. Similar to the aSyn constructs, biotinylated ssDNA was conjugated to the C-terminus via an aldehyde encoding sequence for immobilization and as a reference point. The Bcl-X_L_ protein has a single cysteine that is located close to the C-terminus of the protein (**Fig. 2c**) For the identification of a folded protein with FRET X, an experimentally obtained fingerprint should be mapped against a database consisting of computationally generated protein fingerprints using their online available 3D structures. Hence, we used our previously developed FRET X fingerprint prediction tool that takes into account the effect of the DNA tags on the protein structure^19^ and predicted the fingerprint of Bcl-X_L_ by simulating 200 protein molecules, each being probed 10 times. For the Bcl-X_L_ model protein we predicted that the fingerprint would consist of a single high FRET peak, and this was in line with experimental data (**Fig. 2d,e and Supplementary Fig. 5c**). Taken together, these results show that our FRET X fingerprinting approach is capable of obtaining reproducible fingerprints for both intrinsically disordered and folded human proteins, underscoring that the introduction of additional DNA tags and the labeling procedure itself do not interfere with our fingerprinting approach.

### Post-translational modification mapping

*O*-GlcNAcylation is an essential process in mammalian cells involving the addition of a single N-acetylglucosamine (GlcNAc) to the hydroxyl side chain of serine and threonine residues by *O*-GlcNAc transferase (OGT).^29^ Dysregulation of *O*-GlcNAcylation has been implicated in many human pathologies, such as cancer, diabetes and neurodegenerative diseases, where the PTM site on the protein substrate is decisive for its outcome.^29,30^ However, for *O*-GlcNAcylation, obtaining such information remains challenging, especially as there is no consensus motif for predicting the potential sites of the PTM.^31^ As a result, mapping *O*-GlcNAc sites relies largely on the use of synthetic peptide fragments derived from the protein of interest^32,33^, which may not reflect the *bona fide* PTM sites on the intact protein. Mass spectrometry has been employed to identify sites of *O*-GlcNAcylation. However, it requires proteolytic digestion of intact proteins into peptide fragments, which does not offer the proteoform information. Therefore, due to the lack of methods to characterize the differential enzymatic activity of OGT on distinct Ser/Thr residues on an intact protein, it remains poorly understood how OGT selects Ser/Thr sites to modify. We hypothesized that the high resolving power of FRET X could be leveraged to map potential *O*-GlcNAc sites of a full-length protein.

We incubated the aSyn protein, which is known to undergo *O*-GlcNAcylation with OGT in the presence of uridine diphosphate-linked 6-azido-GlcNAc.^34^ The modified aSyn was subjected to copper click chemistry to attach the donor docking strands to the PTMed residues (**Fig. 3a**) and then immobilized at the C-terminus in a similar fashion as before (**Fig. 1 and Fig. 2**). We observed a main FRET peak with an efficiency of 0.12 and a second FRET peak with an efficiency of 0.23 (**Fig. 3b,c**), indicating that labeling was successful. We compared the FRET efficiencies of the *O*-GlcNAcylated aSyn proteins with those that we obtained for the single cysteine constructs (**Fig. 1**) and found that the FRET efficiency for *O*-GlcNAc modified residues is close to those of aSyn Cys_42_ and Cys_78_, suggesting that the *O*-GlcNAc is attached to residues that are in close proximity (**Fig. 3d**, blue region,). Consistent with these data, mass spectrometry revealed that the *O*-GlcNAcylation had occurred at Thr_54_ and Thr_64_ (**Fig. 3d**, blue spheres, **Supplementary Fig. 6**), underscoring the predictability, accuracy and reproducibility of FRET X and its use for PTM mapping.

**Fig. 3:**
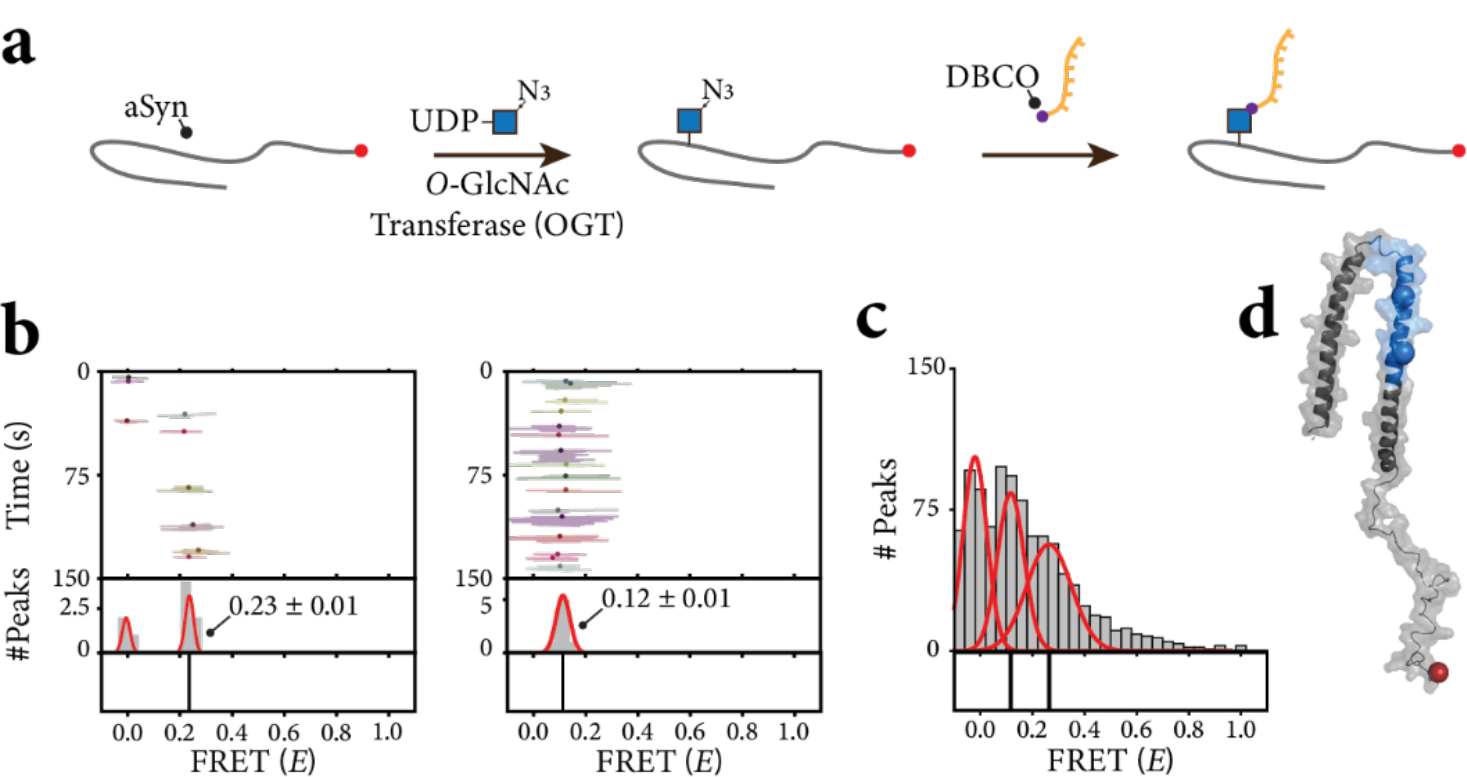
Post-translational modification mapping using FRET X. (**a**) Schematic representation of the PTM labeling scheme. In a first step, the UDP-linked 6-azido-GlcNAc is conjugated to the aSyn substrate via the OGT enzyme. Next, the docking strands are conjugated to the *O*-GlcNAcylated aSyn protein via DBCO click-chemistry. (**b**) Representative kymographs from individual aSyn molecules reporting on the distance of the *O*-GlcNAc to the reference point. The mean FRET ± SEM is reported for each molecule. (**c**) The FRET X histogram and fingerprint for all molecules in a single field of view. We observed two FRET peaks indicating the attachment of two *O*-GlcNAc residues on aSyn, with a FRET efficiency of 0.12 ± 0.12 and 0.25 ± 0.19. These values report on the mean FRET efficiency ± the FWHM of the Gaussian fit. (**d**) Three-dimensional conformation for micelle-bound alpha-synuclein (PDB entry 1XQ8) with the C-terminus shown in red. The proposed region for the PTM sites based on the FRET (*E*) of the cysteines probed in **Fig. 1f** is indicated with the blue shading, while the exact PTM locations are indicated with the blue spheres.

### A universal approach for protein fingerprinting

Finally, we focused on bringing FRET X fingerprinting to natural proteins by circumventing recombinantly expressed tags for immobilization. Such universality is a crucial step towards analyzing specific biomarkers from natural sources.

We developed a labeling approach that allows for the site-selective attachment of a bifunctional linker to the N-terminus of any protein substrate. We made use of the previously reported PCA chemistry^35^ and copper free click chemistry. The modified proteins were subjected to a second labeling step to attach the biotinylated FRET X reference sequence (**Fig. 4a**). To demonstrate site-selective N-terminal modification, we first applied our labeling strategy to our model aSyn protein constructs and as expected we observed a monotonous decrease in FRET efficiency as the distance to the N-terminus increases (**Fig. 4b**, green squares). Furthermore, by combining the FRET fingerprints that were obtained from both N and C-terminus, we were able to probe every region of the full-length aSyn protein (**Fig. 4b**, and **Supplementary Fig. 7**).

**Fig. 4:**
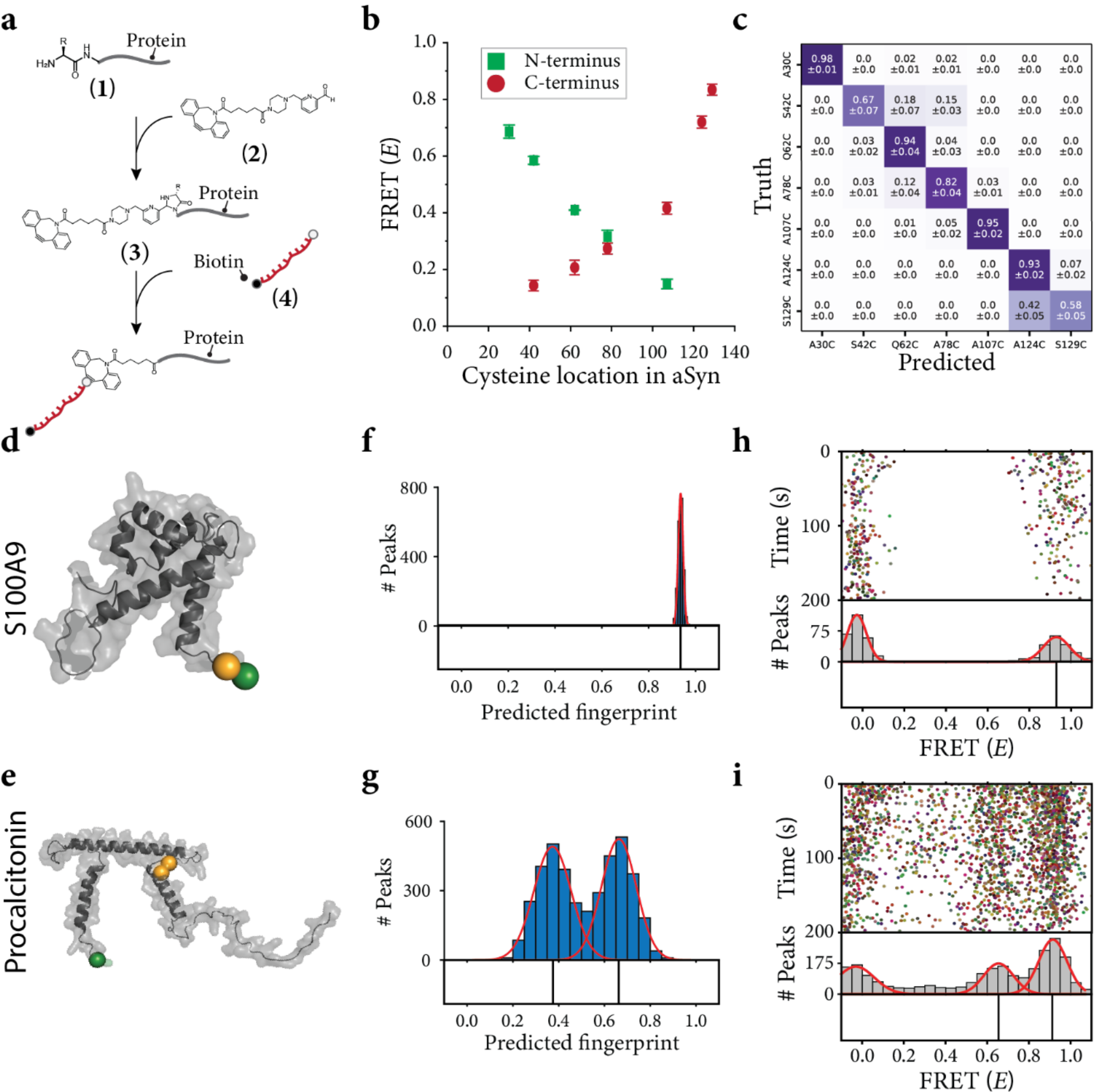
Single-Molecule protein fingerprinting of target proteins using N-terminal labeling. **(a**) Schematic representation of the labeling procedure. The N-terminus of any target protein (1) is labeled with a 2PCA-DBCO derivative (2). The product of this reaction is a protein molecule that has its N-terminus functionalized with a unique DBCO group (3) that allows for the attachment of a biotinylated DNA reference point (4). (**b**) The FRET efficiency as a function of the location of the cysteine in the aSyn protein. A monotonous decrease in FRET efficiency is observed for the cysteine relative to the N-terminus (green squares). The values are reported as the mean ± the standard deviation of three independent experiments. (**c**) Support vector classifier performance on fingerprints for seven aSyn mutants. Heatmaps show how each mutant is classified (mean±standard deviation over 10 cross validation folds), whereby the diagonal positions indicate correct classifications. The N- and C-terminally measured fingerprints were paired to simulate an experiment in which each molecule is sequentially measured from both termini. Classifiers were trained and tested on experimental data from separate experiments, conducted on different days to avoid batch effects. (**d and e**) 3D structures of two inflammatory disease biomarkers S100A9 (**e**, AlphaFold: AF-P06702-F1) and Procalcitonin (PCT) (**e**, AlphaFold: AF-P01258-F1) with the cysteines (orange spheres) and N-terminus highlighted (green sphere). (**f and g)** The predicted FRET fingerprints (mean ± FWHM) for S100A9 protein (f, 0.94 ± 0.02) and PCT (g, 0.37 ± 0.19 and 0.66 ± 0.18). (**h and i**) Experimentally obtained fingerprints reporting on the location of the cysteines to the N-terminal reference point. (**i**) Ensemble kymograph for S100A9 protein with a single high FRET peak (0.93 ± 0.15) and (**j**) for PCT we obtained two high FRET peaks (0.65 ± 0.17 and 0.91 ± 0.16), both mean ± FWHM.

Next, we reasoned that the ability to probe proteins from more than one reference point should increase prediction accuracy. To validate this, we generated a simulated dataset of constructs containing both a C- and N-terminal reference point, by pairing the experimental fingerprints from the C-& N-terminal measurements per construct. We split the data into training and test sets and repeated the classification exercise using a 2-dimensional SVM. This allowed us to classify the seven aSyn constructs with an accuracy of >85% when combining both C and N terminal fingerprints for the same molecule (**Fig. 4c**). This additional information provides a better classification accuracy compared to only C or only N terminus fingerprints (**Supplementary Fig. 7g and f**).

To demonstrate the general applicability of FRET X, we used the N-terminal modification approach on two inflammatory disease biomarkers, the S100A9 protein, informative for severe forms of COVID-19 (**Fig. 4d**)^36^ and Procalcitonin (PCT), used for diagnosis of bacterial infections (**Fig. 4e**)^37^. We ran our fingerprinting simulations and predicted a single high FRET peak for S100A9 protein (**Fig. 4f**), which is in good agreement with our experimental findings (**Fig. 4h**). For PCT we anticipated that resolving the location of both cysteines would be challenging as the difference to the reference point differs by only ∼ 0.2 nm (**Supplementary Fig. 7a**). However, when we predicted the structure, with the DNA labels attached, using our lattice model prediction tool (**Supplementary Fig. 8b,c**), we observed two clearly distinguishable FRET peaks for PCT (**Fig. 4g**). We hypothesize that the larger difference in FRET efficiency between the cysteines is a result from the DNA docking strands, thereby increasing the resolvability. This hypothesis is further supported by the experimental data showing two clear FRET peaks for the PCT biomarker (Cys_85_ and Cys_91_) (**Fig. 4i and Supplementary Fig. 8d,e**). We speculate that the difference in peak position for the predicted and experiment fingerprint is caused by the low prediction power of AlpaFold for the intrinsically disordered regions within PCT (pLDDT <50)^38,39^. The fingerprint simulations are designed to give an indication on the number of FRET and location of peaks in a protein fingerprint and do not always agree with the experimental fingerprints. To demonstrate the fingerprinting ability of FRET X we trained a classifier on experimental data and determined its protein identification accuracy. We observed a mean classification accuracy 80 % (**Supplementary Fig. 8f**) for a protein mixture of Bcl-X_L_, S100A9 and aSyn A124C based on single cysteine fingerprints. This indicates that protein identification is more efficient when a database of experimental fingerprints is constructed and proteins are identified based on this database.

Altogether these results demonstrate the versatile and broad applicability of FRET X to obtain high resolution fingerprints for protein identification and PTM mapping at the single-molecule level.

## Discussion

We introduced a single-molecule fingerprinting approach for protein identification and PTM site mapping. FRET X enables discrimination of proteins with only subtle differences, as well as unambiguous mapping of authentic PTM sites, owing to its ability to pinpoint specific amino acid residues and PTMs on intact proteins. Because the positional information is preserved, structures of the same mass that are generally not discernible in mass spectrometry can be readily differentiated from each other by their distinct FRET efficiencies.

By using short fluorescently labeled DNA strands and their transient binding, FRET X allowed repeated examination of an amino acid residue in a single protein, increasing the localization precision and thereby the overall accuracy of the protein fingerprint.^19,24^ FRET X can fingerprint full-length proteins, such as the intrinsically disordered protein alpha-synuclein and folded proteins such as Bcl-X_L_, S100A9 protein, and PCT. Furthermore, our FRET X fingerprinting approach benefits from the programmable and predictable kinetics of DNA hybridization, which allows for further speed optimization and for multiple target residues to be probed in sequential imaging cycles.^25^ This sequential probing allows us to probe different residues (e.g. amino acids or PTMs) separately by flushing in orthogonal imager strands^24,40^. This strategy avoids crowding of the FRET spectrum, thereby allowing us to resolve the FRET fingerprint for each of the residues with high precision. We have previously shown that by probing cysteines and lysines; or cysteines, lysines and arginines, the uniqueness of a protein fingerprint increases significantly, enhancing the proportion of human proteins that can be identified to 82% or 95%, respectively.^19^

At the current acquisition speed, high-resolution protein fingerprints of several thousand proteins can be obtained within several minutes, which is several orders of magnitude faster than other single-molecule fluorescence protein fingerprinting methods^22^. The capability to fingerprint full-length proteins avoids the need for additional sample preparation steps like digestion into peptides or protein translocation, which are often required for other single molecule protein identification approaches.^12–15^ Since the average protein diameter is estimated to be 5 nm^41^, the typical protein is well within the range of the Cy3-Cy5 FRET pair, while for proteins of different sizes, other FRET pairs may be selected. Furthermore, we have shown that proteins can be immobilized using an N-terminal labeling strategy, thereby removing the need for genetic or synthetic tags, which opens up avenues for the analysis of proteins from natural sources (e.g. body fluids or single cells). While the N-terminus might not always be accessible for labeling^42^, the C-terminus may instead be targeted^43^ for conjugation of the reference and immobilization strand. Additionally, by combining N-, and C-terminus reference points, we are able to expand sequence coverage within a protein (**Figure 4c**).

We further demonstrated that FRET X can be readily exploited to map potential *O*-GlcNAc sites of aSyn. The use of intact proteins is critical in this use case because it better mimics how an enzyme encounters a substrate *in vivo*, as compared to synthetic protein constructs.^29^ Moreover, it also takes into account the crosstalk between *O*-GlcNAc residues at adjacent sites, which is generally neglected when using peptides as substrate. Our results on aSyn *O*-GlcNAcylation were consistent with a previous report in which aSyn expressed in mammalian cells contained up to two *O*-GlcNAc residues^44^. Although we have demonstrated PTM detection using *in vitro* attachment of the modified *O*-GlcNAc, we envision that approach can be further developed for *in vivo* analysis of *O*-GlcNAc-ed protein using metabolic labeling experiments.

Furthermore, FRET X is not limited to mapping *O*-GlcNAcylation sites, but can be readily integrated with existing chemoenzymatic labeling approaches to target acetylation^45^, ribosylation^46^,and fucosylation^47^. Additionally, by combining endoglycosidases and galactosyltransferases, click handles^48,49^ can be incorporated into *N*-glycans to allow for DNA labeling and analysis of full-length glycoproteins. Furthermore, PTM detection using FRET X may go beyond metabolic labeling using other chemical biology strategies to attach orthogonal DNA docking strands to phosphorylation^22,23^ or lipidation^50^.

One of the main challenges for high-throughput single-molecule proteomics lies in the varying abundance of different protein species in the cell. The dynamic range of the proteome spans several orders of magnitudes^1,51^, due to which low abundant species can easily get masked by more abundant ones. Owing to its single-molecule sensitivity and the ability to fingerprint several thousands of proteins in a single field of view, our method detects even the sparsest proteins, and future optimizations including automated acquisition and scanning stages might increase throughput and thereby sensitivity even further. Alternatively, we may address the challenge posed by the large dynamic range by adopting protein enrichment strategies for a targeted approach.^52^ In the current study, less than a femtomole of labeled protein was needed for fingerprinting and when improvements, such as automated microfluidics, can be incorporated in our assay, the sensitivity may be further increased with one or two orders of magnitude.

To conclude, we demonstrated a single-molecule protein fingerprinting approach that allows for localization of both amino acids and PTMs in full-length proteins. We envision that our full-length single-molecule protein fingerprinting may provide a tool for proteomics at the ultimate sensitivity.

## Author Contributions

M.F. and C.J. initiated and designed the project. M.F. designed and performed the protein labeling procedures. M.F. and R.W. performed the single-molecule FRET X experiments. I.W. and C.A.P. expressed and purified the proteins. C.L. wrote the software for and performed the fingerprinting predictions and protein classification. D.R. supervised the fingerprinting prediction simulations. S.H.K. wrote the automated peak finding code for single molecule FRET X analysis. M.F. and Z.L. conceptualized the *O*-GlcNAc site mapping and designed the chemoenzymatic strategy. M.F. and Z.L. designed the N-terminal labeling strategy. Y.F. and G.-J.B. synthesized UDP-Azido-*O*-GlcNAc for protein *O*-GlcNAcylation. M.F., S.H.K., C.d.L., and C.J. analyzed and discussed the data. M.F., R.W., and C.J. prepared the initial draft of the manuscript. All authors read and approved the manuscript.

## Acknowledgements

C.J. and D. R. acknowledge funding from NWO-I (SMPS). C.J. acknowledges funding from NWO (Vici), Basic Science Research Program (NRF), and Frontier 10-10 (Ewha Womans University). Y.F. was funded by the Chinese Scholarship Council (CSC) Grant.

## Declaration of interests

C.J., M.F., C.L., and D.R. hold a patent on single-molecule FRET for protein characterization (patent number: WO2021049940). C.J., M.F., Z.L. filed a patent for the bifunctional linker for N-terminus protein modification.

## Data availability

The data supporting the main finding of this study are available in the article and its supplementary information. Any additional data are available from the corresponding author upon request.

## Code availability

The algorithms for the codes supporting the main findings of this study are available in the Article and its Supporting information. Any additional information concerning the code is available from the corresponding author upon request.

