## Supplementary Information for "Full-Length Single-Molecule Protein Fingerprinting"

**This PDF file includes:**

Figures. S1 to S7

Tables S1 to S2

Materials and Methods

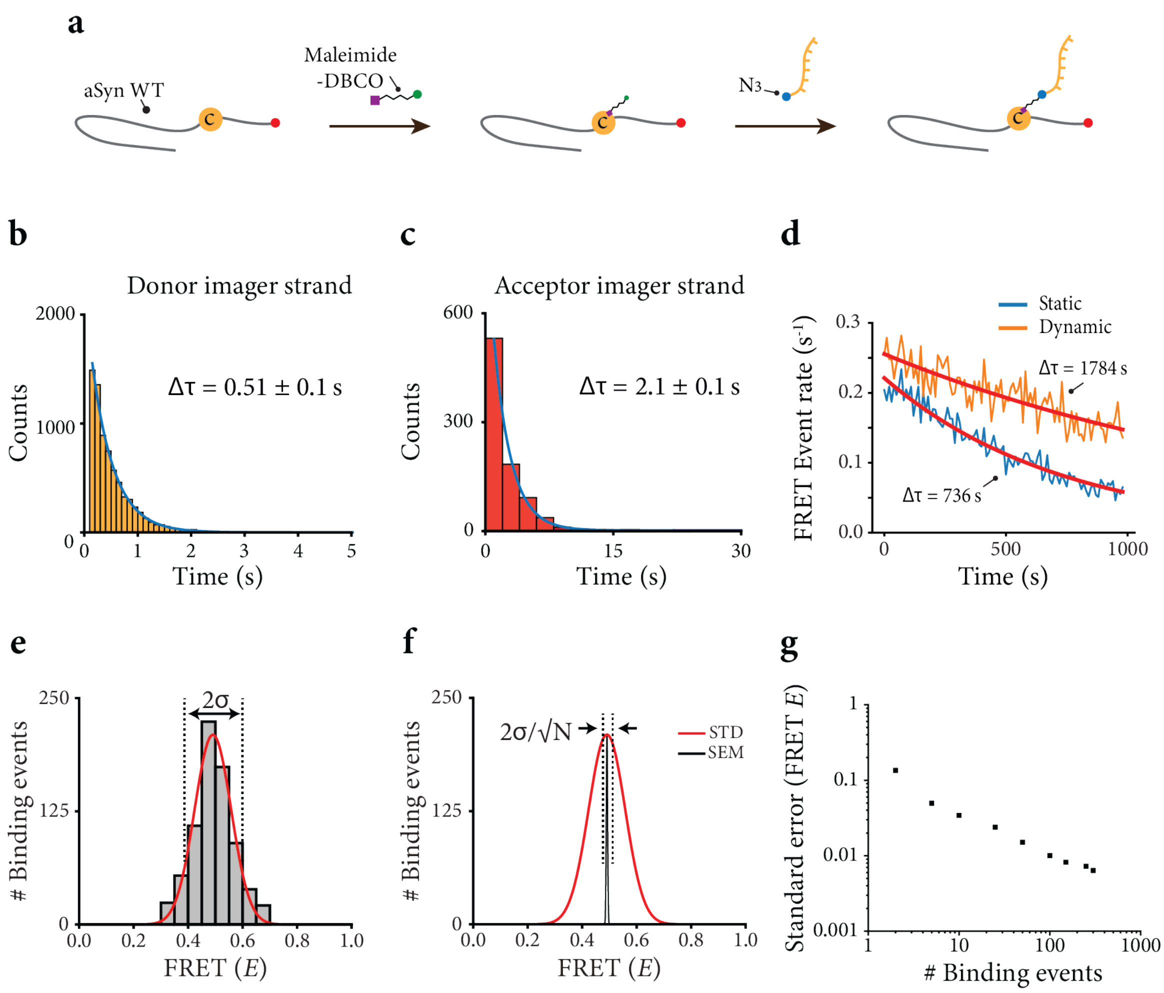


**Fig. S1: General approach for FRET X fingerprinting.** (**a**) Schematic representation of the cysteine labeling scheme for aSyn and other proteins used in the study. (**b**) Dwell-time histogram for the FRET X donor imager strand (Table S2) fitted with a maximum likelihood estimation for a single exponential distribution (blue line) (n=4288). Average ± standard deviation of four different estimates gives: 0.51 ± 0.1 s. Protein substrate was aSyn$_{Cys78}$. (**c**) Dwell-time histogram for the FRET X acceptor imager strand (Table S2) fitted with a maximum likelihood estimation for a single exponential distribution (blue line) (n=870). Average ± standard deviation of four different estimations gives: 2.1 ± 0.1 s. Protein substrate was aSyn$_{Cys78}$. (**d**) The time-dependant FRET detection rates for a static acceptor vs a dynamic acceptor were calculated with time windows of 20 s from 500 aSyn molecules under constant illumination. The static acceptor showed fasted decrease in the FRET event rate due to photobleaching. (**e**)The standard deviation (σ) reports on the intrinsic broadness of a FRET histogram. (**f**) The standard error measures the accuracy of determining the centre of a peak. N is the number of imager strand binding events. The analysis of panels **d** and **e** was performed on aSyn$_{Cys107}$. (**g**) Standard error of the FRET X efficiency versus the number of binding events. We observe that we can determine the center of a Gaussian fit with a FRET X precision of $\Delta E \sim 0.03$ after >10 binding events.

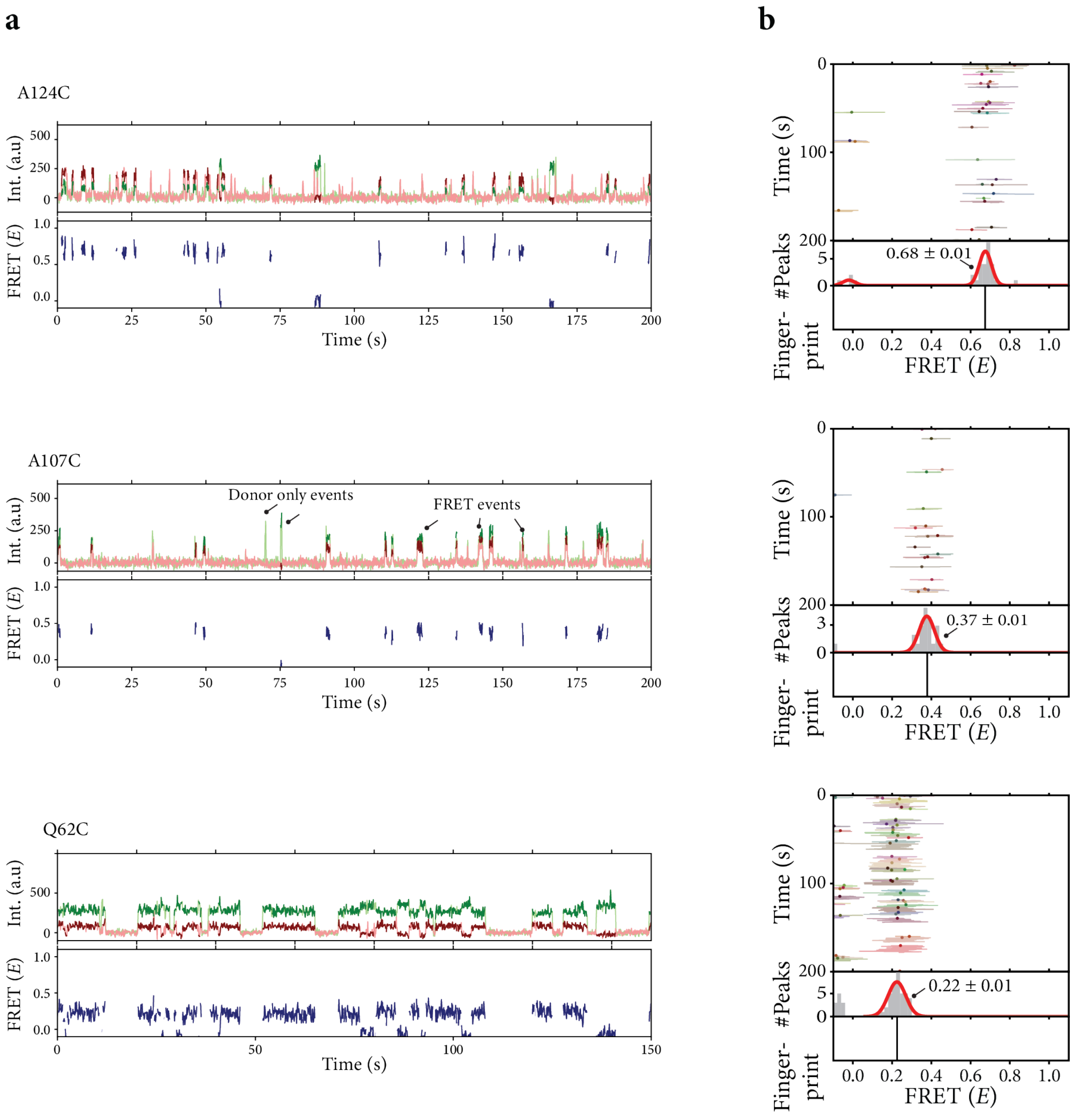


**Fig. S2: Representative single molecule time traces and kymographs.**

(**a**) Example single-molecule fluorescence and FRET time traces obtained for different aSyn constructs. For each of the constructs we observed repetitive short lived binding events for long periods of time. (**b**) The single molecule kymographs are constructed from each individual time trace, where the lines indicate the FRET efficiency per data point within a FRET events and the dots are the mean FRET efficiency per event. The mean FRET efficiencies were fitted with a Gaussian and the FRET efficiency is reported as the mean ± SEM for the Gaussian fits.

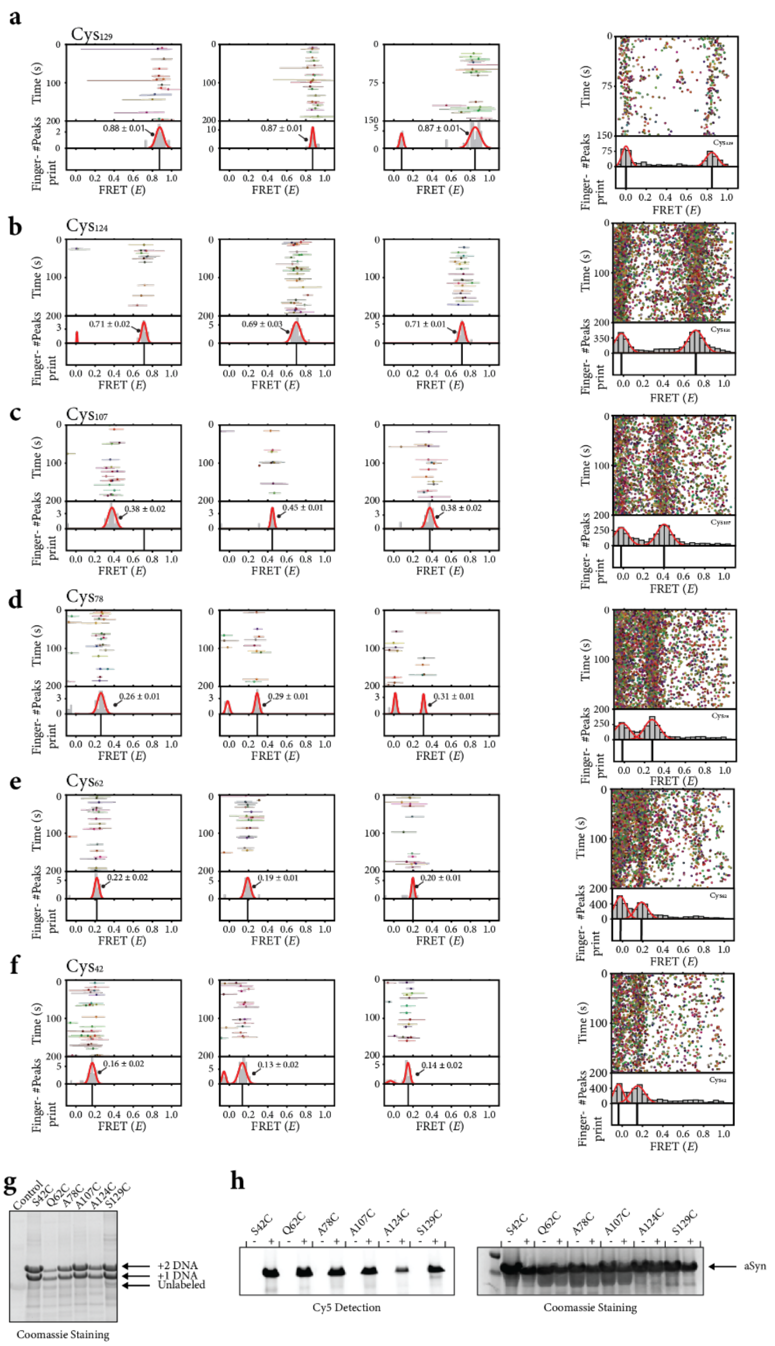


**Fig. S3: Representative FRET kymographs, histograms and fingerprints for single cysteine mutants of alpha-synuclein probed from the C-terminus.**

(**a-f**) Single-molecule (first 3 panels) and ensemble (right) FRET kymographs obtained for each of the different aSyn mutants measured in **Fig. 1. a-f**) Reported values in the single-molecule kymographs are mean ± SEM for the Gaussian fits. For the ensemble kymographs we observed FRET efficiencies of 0.83 ± 0.13 for Cys129, 0.71 ± 0.15 for Cys124, 0.40 ± 0.17 for Cys107, 0.27 ± 0.17 for Cys78, 0.19 ± 0.14 for Cys62, and 0.15 ± 0.13 for Cys42 (all mean ± FWHM of the Gaussian fits). (**g**) The cysteine labeling efficiency was determined by SDS-PAGE gel electrophoreses for aSyn double cysteine constructs and Bcl-$X_L$. We note that for these FGE-recombinantly expressed proteins we observed an extra DNA band on the gel ((#Cys in protein) + 1). This is the result from unsuccessful conversion of the FGE-tag to a formyl-glycine (aldehyde) leaving an extra cysteine that can be labelled with DNA. These protein can not be labeled with a hydrazide-DNA-biotin strand and will therefore not be immobilised on the surface. The labeling efficiency for S42C was 45% for one DNA and 55% for two DNA, for Q62C was 53% for one DNA, and 40% for two DNA, for A78C was 48% for one DNA and 49% for two DNA, for A107C was 45% for one DNA and 53% for two DNA, for A124C was 52% for one DNA and 44% for two DNA, and for S129C was 52% for one DNA and 43% for two DNA. (**h**)The FGE labeling efficiency was low and thus difficult to visualize directly with DNA conjugation, therefore we conjugated a fluorophore directly to the FGE tag. The labeling efficiency was determined by measuring the absorbance of protein and Cy5 using nanodrop and the efficiency was 8% for S42C, 9% for Q62C, 7% for A78C, 7% for A107C, 4% for A124C, and 6% for S129C.

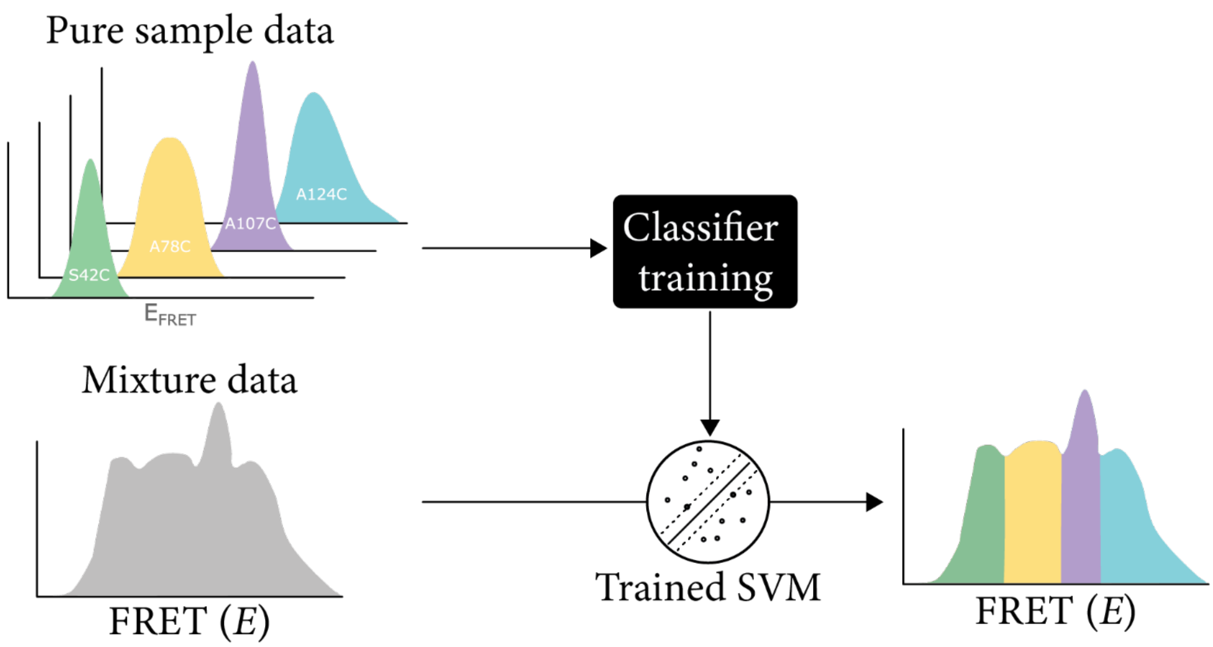


**Fig. S4**: Schematic overview of the approach used to separate the FRET spectrum in different regions to classify alpha synuclein mutants in a mixture. A support vector machine (SVM) classifier was first trained on FRET values obtained from four separate experiments each containing a single aSyn mutant to identify the FRET region associated with each mutant. The trained SVM was then used to classify the mutants based on their FRET values in a mixture and to determine the relative concentrations of each of the constructs.

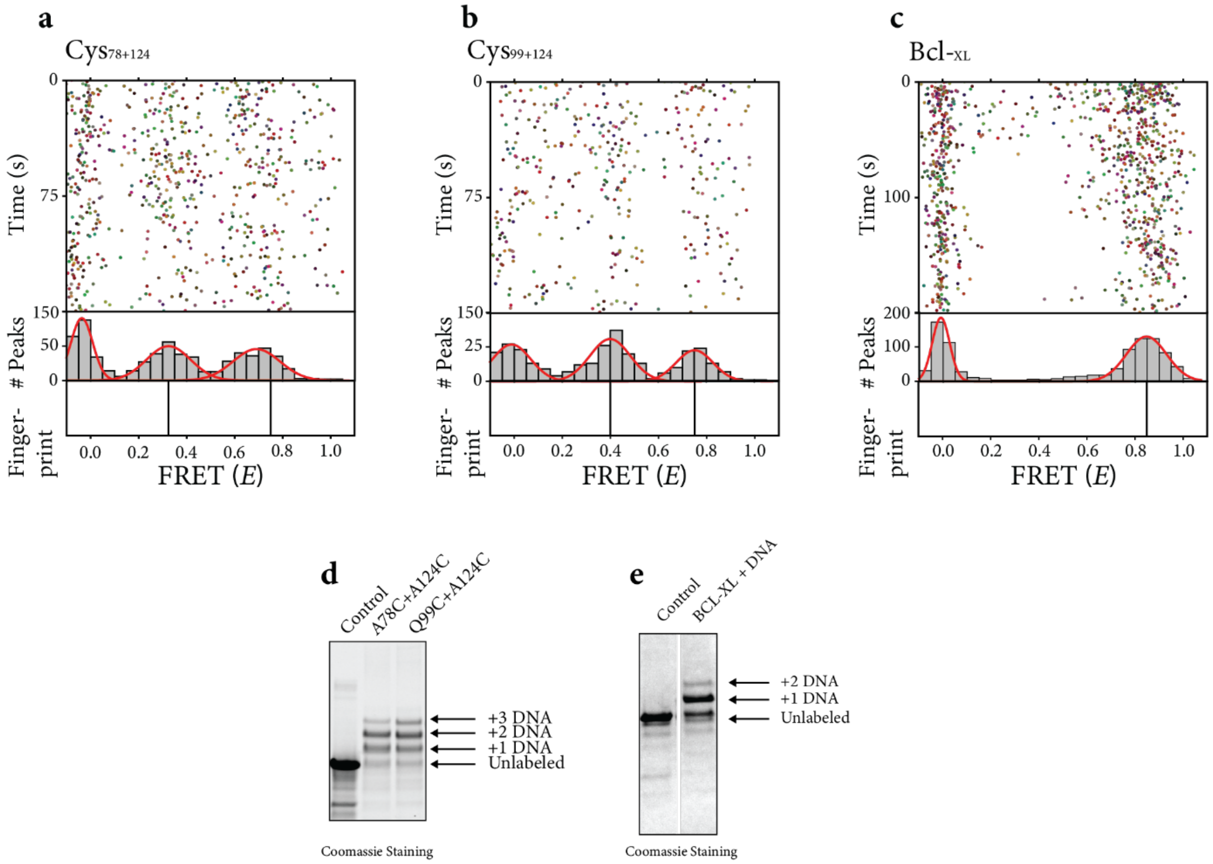


**Figure S5: Representative FRET kymographs for biomarkers probed from the C-terminus.**
(**a-c**) Ensemble FRET kymographs obtained for each of the distributions shown in **Figure 2**. For aSyn-$Cys_{78}$+$Cys_{124}$ (**a**) are 0.29 ± 0.19 and 0.69 ± 0.10 and for aSyn-$Cys_{99}$+$Cys_{124}$ (**b**) 0.43 ± 0.14 and 0.77 ± 0.16. (**c**) For Bcl-$X_L$ we observed a FRET efficiency of 0.85 ± 0.20. All FRET efficiencies are reported as the mean ± FWHM of the Gaussian fit. (**d and e**) The cysteine labeling efficiency was determined by SDS-PAGE gel electrophoreses for aSyn double cysteine constructs and Bcl-$X_L$. We note that for these FGE-recombinantly expressed proteins we observed an extra DNA band on the gel ((#Cys in protein) + 1). This is the result from unsuccessful conversion of the FGE-tag to a formyl-glycine (aldehyde) leaving an extra cysteine that can be labelled with DNA. These protein can not be labeled with a hydrazide-DNA-biotin strand and can therefore not be immobilised on the surface. The labeling efficiency for $aSyn^{A78C+A124C}$ was 35% for one DNA, 42% for two DNA, and 8% for 3 DNA, for $aSyn^{Q99C+A124C}$ was 30% for one DNA, 40% for two DNA and 15% for 3 DNA. The Bcl-$X_L$ labeling efficiency was 70% for one DNA and 8% for two DNA.

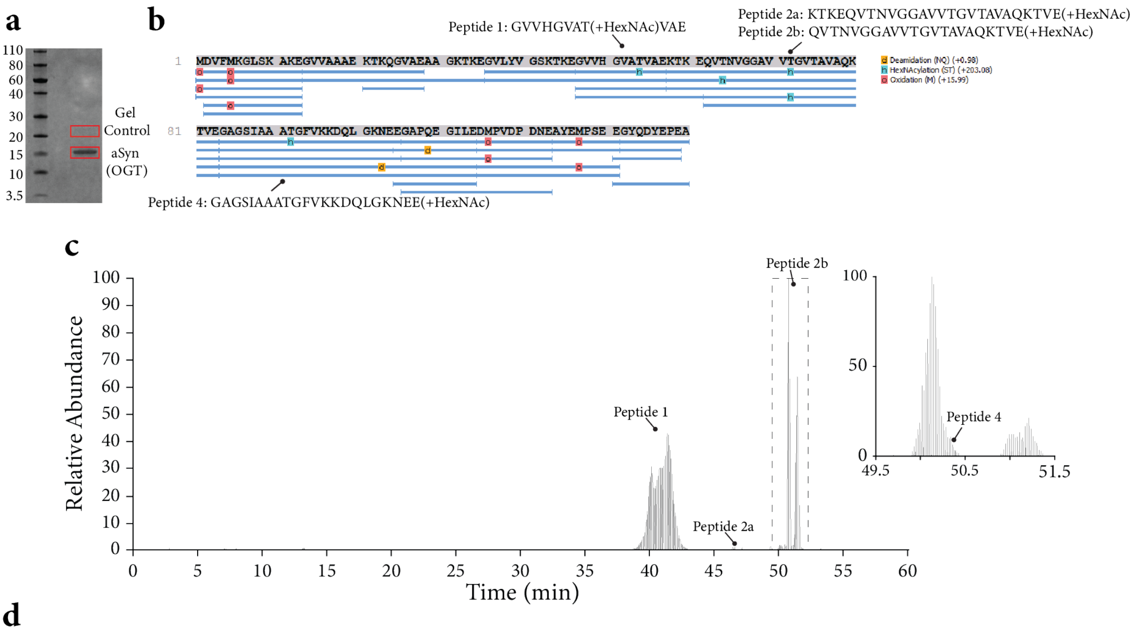


**d**

Gel Control

| Protein Group | Protein ID | Accession | -10lgP | Coverage (%) | Coverage (%) Sample 7 | Area Sample 7 | #Peptides | #Unique | #Spec Sample 7 | PTM | Avg. Mass | Description |
|---|---|---|---|---|---|---|---|---|---|---|---|---|
| total 0 proteins | | | | | | | | | | | | |

aSyn (OGT)

| Protein Group | Protein ID | Accession | -10lgP | Coverage (%) | Coverage (%) Sample 8 | Area Sample 8 | #Peptides | #Unique | #Spec Sample 8 | PTM | Avg. Mass | Description |
|---|---|---|---|---|---|---|---|---|---|---|---|---|
| 1 | 1 | aSyn_com | 164.30 | 100 | 100 | 8.18E8 | 15 | 15 | 90 | Y | 14460 | aSyn_com |
| total 1 proteins | | | | | | | | | | | | |

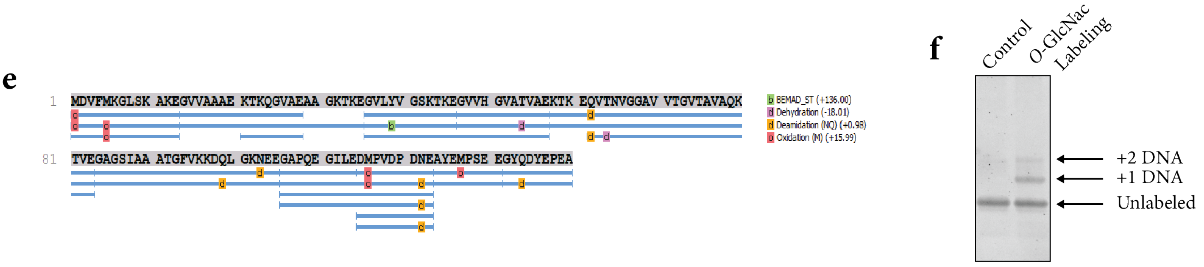


**Fig. S6: Degree of HexNAcylation of aSyn and identification of glycosylated amino acid residues by glycoproteomics.** (**a**) Coomassie stained gel reporting on which regions were extracted from the gel for the negative control and the O-GlcNAcylated aSyn protein, (**b**) Obtained sequence coverage following GluC in-gel digestion and (glyco)proteomic analysis of the resulting peptides. Thereby, three peptides showed additional HexNAcylated peptide variants. (**c**) Extracted ion chromatogram for the HexNAcylated forms of Peptide 1 (621.3156-621.3296), Peptide 2b (705.1279-705.1470) and Peptide 3 (798.7357-798.7521). Peptide 1 and Peptide 2 (a+b) showed a very abundant peak for the modified peptide variants. However, the signal for the modified variant of Peptide 3 was comparability low abundant. (**d,e**) No peptides related to a-syn were detected in the analysed gel control. Furthermore, beta-elimination on the in-gel digested peptides was performed to eliminate the HexNAc modifications. This results in a modified serine/threonine (dehydration) in the peptide which can be detected in the proteomics experiment. Ultimately, this allowed to identify two threonine residues in peptide 1 (ATV) and peptide 2 (VTN) as HexNacylated residues. (**f**) The *O*-GlcNac labeling efficiency was determined by SDS-PAGE gel electrophoreses for aSyn. We observed two additional bands, that suggests attachment of DNA to two different *O*-GlcNAc. The labeling efficiency for one DNA was 30 % and for two DNA was 15 %.

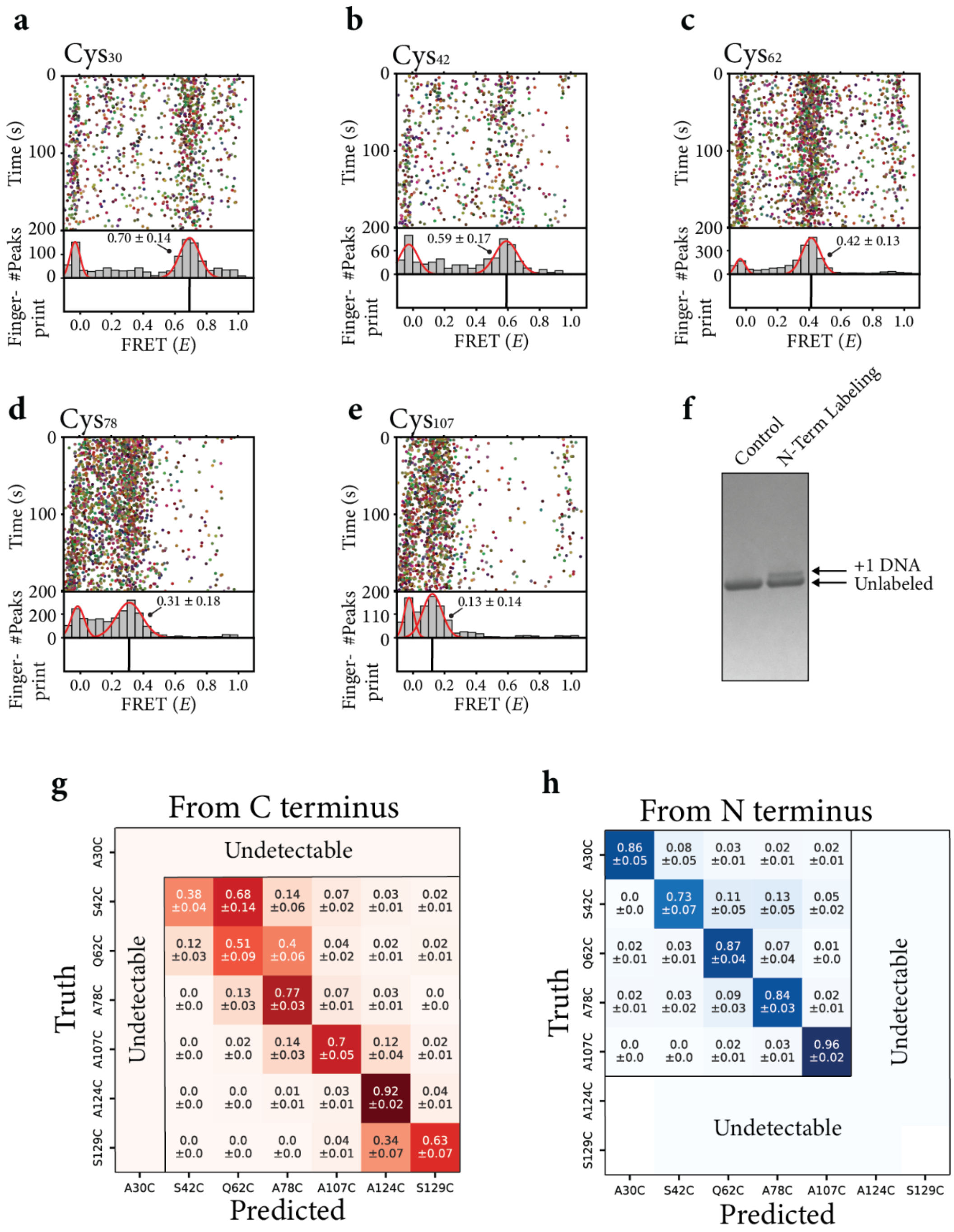


**Fig. S7: Representative FRET kymographs, histograms and fingerprints for single cysteine mutants of alpha-synuclein probed from the N-terminus.**

(**a-e**) Single-molecule ensemble FRET kymographs obtained for each of the different aSyn mutants measured in **Fig. 4 a-e**). Reported values in the kymographs are mean ± FWHM of the Gaussian fits. (**f**) The N terminal labeling efficiency was determined by SDS-PAGE gel electrophoreses for aSyn. We observed a single additional band, confirming that only one modification had occurred. The overall labeling efficiency was 40 %. (**g and h**) Confusion matrix to determine classification accuracy for each aSyn construct measured from C terminus (g) or N terminus (h). For each of the mutants, experimentally determined FRET X values were used to train a support vector machine (SVM) classifier. Trained SVMs were subsequently evaluated on FRET X values obtained in different experiments. Note that some cysteines are undetectable from the N or C terminus. The procedure was repeated 100 times on bootstrapped training and test data to obtain confidence intervals around classifier performance measures.

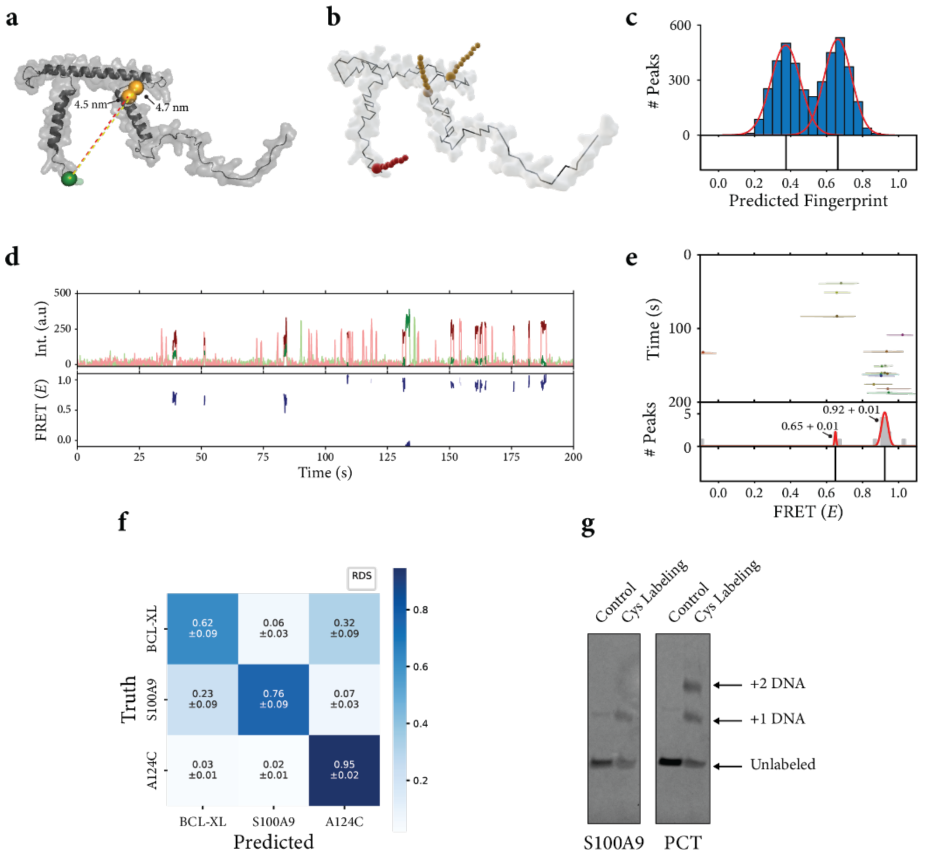


**Fig. S8: Schematic of FRET X fingerprinting simulation for Procalcitonin.**

(**a**) 3D structure of Procalcitonin (PCT) (AlphaFold: AF-P01258-F1) and the distance of the cysteines to the N-terminal reference point. (**b**) The PCT protein structure is loaded into our lattice model. In the lattice model, all residues are reduced to the Cα positions. The residues to which the DNA is attached are highlighted (orange spheres for cysteines, red spheres for N-terminus), after which the structure is randomly mutated using a Markov Chain Monte Carlo (MCMC) process, until docking strands no longer experience steric hindrance from the rest of the protein structure.[19] (**c**) The MCMC process continues while snapshots are taken at regular intervals. The distances for each FRET pair are  averaged over all snapshots and translated into a FRET efficiency to produce the FRET fingerprint for this protein. (**f**) a three-way confusion matrix that represents the mean classification accuracy for each of the biomarkers. For the protein mixture, experimentally determined FRET X values were used to train a support vector machine (SVM) classifier. Trained SVMs were subsequently evaluated on FRET X values obtained in different experiments. The mean classifier accuracy for the three biomarkers was 80% based on single cysteine protein fingerprints. (**g**) The cysteine labeling efficiency was determined by SDS-PAGE gel electrophoreses for biomarkers S100A9 and PCT. The gel was Coomassie stained and an extra band was observed for S100A9, repressing attachment of a single DNA (efficiency 65%) and we observed two additional bands for PCT, representing two cysteines labeled with DNA(the labeling efficiency one DNA was 40 % and for two DNA 31 %).

**Supplementary Table 1: Protein sequences constructs**

| Protein name | Sequence (N → C) | Supplier |
|---|---|---|
| Alpha-Synuclein | MDVFMKGLSKAKEGVVAAAEKTKQGVAEAAGKTKEGVLYVGSKTKEGVVHGVATVAEKTKEQVT NVGGAVVTGVTAVAQKTVEGAGSIAAATGFVKKDQLGKNEEGAPQEGILEDMPVDPDNEAYEMPSE EGYQDYEPEALCTPSRYQDPVQVDAAAELALVPRGSSAHHHHHHHHHH | Genscript |
| BCL-XL | MGSSHHHHHHSSGLVPRGSHMSQSNRELVVDFLSYKLSQKGYSWSQFSDVEENRTEAPEGT ESEMETPSAINGNPSWHLADSPAVNGATGHSSSLDAREVIPMAAVKQALREAGDEFELRYRRAFSDLT SQLHITPGTAYQSFEQVVNELFRDGVNWGRIVAFFSFGGAL**C**VESVDKEMQVLVSRIAAWMATYLN DHLEPWIQENGGWDTFVELYGNNAAAESRKGQERFNRWFLTGMTVAGVVLLGS LFSRKLCTPSR | Genscript |
| Procalcitonin | MGFQKFSPFL ALSILVLLQA GSLHAAPFRS ALESSPADPA TLSEDEARLL LAALVQDYVQ MKASELEQEQ EREGSSLDSP RSKR**C**GNLST**C**MLGTYTQDF NKFHTFPQTA IGVGAPGKKR DMSSDLERDH RPHVSMPQNA N | Bio-Techne |
| S100A9 | MT**C**KMSQLER NIETIINTFH QYSVKLGHPD TLNQGEFKEL VRKDLQNFLK KENKNEKVIE HIMEDLDTNA DKQLSFEEFI MLMARLTWAS HEKMHEGDEG PGHHHKPGLG EGTP | Aviva System Biology |

**Supplementary Table 2: DNA constructs**

| DNA Strand | Sequence (5’ - 3’) | Modification | Supplier |
|---|---|---|---|
| Donor imager strand | CTCCTC | 3’ Cy3 | Ella Biotech (GmbH) |
| Acceptor imager strand | TAATGAAGA | 3’ Cy5 | Ella Biotech (GmbH) |
| Donor docking sequence - Cysteine | TATACATCTAT | 5’ Azidobenzoate | Biomers.net (GmbH) |
| Donor docking sequence – O-GlcNAc | TATACATCTAT | 5’ DBCO | Biomers.net (GmbH) |
| Acceptor docking sequence – FGE | TTCTTCATTACTATTTTTTTTT | 5’ Hydrazide, 3’ Biotin | Biomers.net (GmbH) |
| Acceptor docking sequence – N-terminus | TTCTTCATTACTATTTTTTTTT | 5’ Hydrazide, 3’ Biotin | Biomers.net (GmbH) |

### Materials and Methods

#### Protein expression and purification.

All proteins were codon optimized for *E. coli* BL21 (DE3) and inserted into a pET52b (+) for alpha-synuclein or pET15b for BCL2L1-$X_L$ (see **Supplementary Table 1** for full list of protein sequences). All proteins were engineered to contain an C-terminal aldehyde encoding sequence. The cysteine in this motif is converted *in vivo* into formylglycine by co-expression of the formylglycine-generating enzyme (FGE)[26]. The plasmids and protein encoding genes were synthesized and prepared by GenScript.

The proteins were expressed in *E. coli* BL21(DE3) cells. Cultures were grown at 37 °C in LB medium supplemented with 50 µg/mL kanamycin and 50 µg/mL ampicillin until an $OD_{600}$ of 0.5 was reached. The expression of FGE was induced through addition of 1% L-arabinose at 37 °C and after 30 minutes the expression of the model proteins was induced by 1 mM isopropyl-β-D-thiogalactopyranoside (IPTG). The cultures were transferred to 26 °C to allow for expression of the proteins for 5 hours, after which the cells were harvested at 4,000g. The cells were lysed by resuspending the pellet in 10 mL lysis buffer (50 mM HEPES-KOH pH 7.5, 500 mM NaCl, 0.5% Triton X-100). For the alpha-synuclein proteins, the cells were lysed by boiling the cell suspension for 15 minutes. The cells containing BCL2L1-$X_L$ proteins were lysed by tumbling the cell suspension for 2 hours at room temperature, followed by sonication on ice during 6 cycles of 30 s ON and 1 min OFF at 30% amplitude. Next, the cell lysate of each model protein was centrifuged at 30,000g for 30 minutes at 4 °C. The proteins were purified from the cell-free extract using HisPur™ Ni-NTA resin according to the manufacturer's manual and buffer exchanged into storage buffer (50 mM HEPES-KOH pH 7.5, 150 mM NaCl, 10% glycerol, 25 mM TCEP) using 10 kDa Amicon Ultra centrifugal filters. All proteins were aliquoted and stored at -80 °C

#### Cysteine labeling.

After purification, proteins were labeled without chemical, temperature or mechanical denaturation to preserve their structure. First, cysteine residues were reduced with 40-fold molar excess (TCEP) for 30 minutes and then labeled with 25-fold molar excess monoreactive maleimide- DBCO in 50 mM HEPES pH 7.5, 150 mM NaCl 1% Triton X-100 buffer overnight at room temperature (23 ± 1°C). Excess maleimide-DBCO and TCEP was removed with Zeba™ Spin desalting columns 7kDa MWCO (ThermoFisher) and the reaction buffer was changed into 0.3 M NaAc pH 5.5, for the aSyn proteins and 50 mM HEPES pH

6.9, 150 mM NaCl, 1% Triton X-100, for Bcl-$X_L$. Then monoreactive Azidobenzoate-(5') functionalized DNA was added in 10-fold molar excess and incubated overnight at room temperature. The formylglycine residues were acceptor-labeled with 10-fold excess biotinylated and hydrazide-functionalized DNA for 46 hours at room temperature for aSyn and 96 hours at 4 °C for Bcl-$X_L$ in a rotary shaker. Free hydrazide-DNA-biotin was removed with Ni-NTA Magnetic Agarose Beads (Qiagen) according to manufacturer's protocol. See **Supplementary Table 2** for the full list of substrates.

***O*-GlcNAcylation labeling.**

The alpha-Synuclein constructs were *O*-GlcNAcylated using a recombinant human *O*-GlcNAcTransferase (OGT) (Novus Biologicals). The reaction was performed by adding 20 excess of aSyn over OGT in a reaction buffer (25 mM Tris pH 7.5, 10 mM $CaCl_2$ and 10 mM $MgCl_2$) supplemented with 10 mM UDP-Azido-*O*-GlcNAc, and incubated overnight at 37 °C. the next day, excess UDP-Azido-*O*-GlcNAc was removed with Zeba™ Spin desalting columns and the reaction buffer was exchanged into 0.3 M NaAc pH 5.5. The *O*-GlcNAc residues were labeled with 10-fold excess DBCO functionalized DNA and the formylglycine residue was labeled with 10-fold excess biotinylated and hydrazide-functionalized DNA for 48 hours at room temperature (23 ± 1 °C) in a rotary shaker. Free DNA was removed with Ni-NTA Magnetic Agarose Beads (Qiagen) according to the manufacturer's protocol.

**N-terminal modification.**

For the N-terminal labeling of protein substrates, we made a 2PCA-DBCO bifunctional linker. For this, we incubated 100 mM of a 2PCA intermediate (6-(piperazin-1-ylmethyl)-2-pyridinecarboxaldehyde HCl salt, (Sigma Aldrich: 808571) with 2-fold excess NHS-DBCO and 3-fold excess of triethylamine in DMSO for 24 hours at room temperature, while shaking. Next, we quenched the NHS by adding 10-fold excess dimethylamine and incubated for 4 to 6 hours at room temperature. The reaction mixture was dried using speed vac and dissolved in DMSO to a concentration of 100 mM 2PCA-DBCO.

The target proteins (alpha-synuclein, Human recombinant Procalcitonin (Bio-Techne) and Human recombinant S100A9 protein (NovusBio) were dissolved in PBS at a concentration of 5 μM. To this, we added 400-fold excess of the 2PCA-DBCO linker and incubated for 24 hours at 37 °C while shaking. The next day, free 2PCA-DBCO was removed with Zeba™ Spin desalting columns 7kDa MWCO. Next the proteins were labeled with 2-fold excess biotinylated and Azide-functionalized DNA for 48 hours at room temperature (23

± 1 °C) in a rotary shaker. Free DNA was removed with Ni-NTA Magnetic Agarose Beads (Qiagen) according to the manufacturer's protocol. Finally, the eluted proteins were reduced with 40-fold molar excess TCEP for 30 minutes and then labeled with 20-fold molar excess monoreactive maleimide-DNA for 24 hours.

**Single-molecule setup.**

All experiments were performed on a custom-built microscope setup. An inverted microscope (IX73, Olympus) with prism-based total internal reflection was used. In combination with a 532 nm diode-pumped solid-state laser (Compass 215M/50mW, Coherent). A 60x water immersion objective (UPLSAPO60XW, Olympus) was used for the collection of photons from the Cy3 and Cy5 dyes on the surface, after which a 532 nm long pass filter (LDP01-532RU-25, Semrock) blocks the excitation light. A dichroic mirror (635 dcxr, Chroma) separates the fluorescence signal which is then projected onto an EM-CCD camera (iXon Ultra, DU-897U-CS0-# BV, Andor Technology). Our pixel size is 107 x 107 nm and the complete field of view is 512x256 pixels (54.8 µm x 27.4 µm) and contains ± 200 molecules. A series of EM-CDD images was recorded using custom-made program in Visual C++ (Microsoft).

**Single-molecule data acquisition.**

Single-molecule flow cells were prepared as previously described[51]. In brief, to avoid non-specific binding, quartz slides (G. Finkerbeiner Inc) were acidic piranha etched and passivated twice with polyethylene glycol (PEG). The first round of PEGylation was performed with mPEG-SVA (Laysan Bio) and PEG-biotin (Laysan Bio), followed by a second round of PEGylation with MS(PEG)4 (ThermoFisher). After assembly of a microfluidic chamber, the slides were incubated with 20 µL of 0.1 mg/mL streptavidin (Thermofisher) for 2 minutes. Excess streptavidin was removed with 100 µL T50 (50mM Tris-HCl, pH 8.0, 50 mM NaCl). Next, 50 µL of 75 pM DNA-labeled protein was added to the microfluidic chamber. After 2 minutes of incubation, unbound protein was washed away with 200 µL T50. Then, 50 µL of 10 nM donor labeled imager strands and 50 nM acceptor labeled imager strands in imaging buffer (50 mM Tris-HCl, pH 8.0, 500 mM NaCl, 0.8 % glucose, 0.5 mg/mL glucose oxidase (Sigma), 85 ug/mL catalase (Merck) and 1 mM 6-hydroxy-2,5,7,8-tetramethylchroman-2-carboxylic acid (Trolox) (Sigma)) was injected. All single-molecule FRET experiments were performed at room temperature (23 ± 1 °C). See Table S.2 for the full list of docking and imager strands.

**Data analysis.**

Fluorescence signals are collected at 0.1-s exposure time. During the acquisition of the movie, the green laser is used to excite the Cy3 donor fluorophores. The fluorescence images were analysed by a custom script written in IDL. The script collects the individual intensity hotspots in the acceptor channel and pairs them with intensity hotspots in the donor channel, after which the time traces are extracted. The details of the automated detection of individual imager strand binding events from the fluorescence time traces are described elsewhere[23,43]. Briefly, a two-state K-means clustering algorithm was applied on the sum of the donor and acceptor fluorescence intensities of individual molecules to find an intensity threshold, with which the trace were divided into high- or low-intensity segments. The high-intensity segments that lasted for more than three consecutive frames were selected for further analysis. If there were abrupt donor and acceptor intensity changes within a high-intensity segment possibly due to photo-bleaching or imager dissociation, the data points that come after the transition moment were removed from the segment. The gamma and leakage factors to were determined from acceptor bleaching events and donor only events, respectively.[44] Average FRET efficiencies from each selected segment were used to build the FRET kymograph and histogram. Populations in the FRET histogram are automatically classified using a Gaussian mixture model (GMM). GMMs containing one to five distributions are fitted, after which the best fitting GMM is selected using the Bayesian information criterion (BIC). Peaks with weights lower than 0.2 are discarded, as these were found to capture background noise. The automated analysis code in Python is freely available at: https://github.com/kahutia/transient_FRET_analyzer2.

**Protein Fingerprint Simulation.**

The protein fingerprinting simulations were performed as previously described.[19] A protein folding simulation was implemented to incorporate DNA-tags attached to certain residues and account for their effect on the protein structure. Lattice models were used because of the far lower computational power needed for folding simulations compared to fully atomistic models that allow unrestricted movement. The lattice model is obtained by reducing each amino acid to a pseudo-atom and restricting its possible positions to the vertices of a body-centered cubic lattice. Such models have previously been used in applications where low computational requirements were essential. The procedure starts with a fully atomistic native structure predicted by AlphaFold2[36,37], which is converted to a lattice structure with tagged residues marked. This structure is assigned a modeling energy $E_{tot}$, based on interactions between pseudo-atoms located on adjacent vertices, the presence of native secondary structures and steric hindrance between pseudo-atoms and DNA-tags:

$$E_{tot} = E_{AA} + E_{sol} + E_{ss} + E_{dsb} + E_{tag} + E_{reg}$$

Here $E_{AA}$ and $E_{sol}$ represent the sums of pairwise residue interaction energies and residue-solvent interaction energies respectively. $E_{ss}$ rewards formation of native secondary structures and $E_{dsb}$ rewards disulfide bridges. $E_{tag}$penalizes steric hindrance between DNA-tags and other pseudo-atoms. Finally $E_{reg}$ penalizes large single-step changes in structure to better retain overall structure. $E_{tot}$ is then minimized using a Markov chain Monte Carlo (MCMC) process, by repeatedly applying random perturbations to the structure and accepting or rejecting them based on the incurred change in the model energy. Further MCMC iterations are used to generate hundreds of slightly different structures, from which distances between donor and acceptor dye positions are deduced. These values are then translated to FRET efficiencies $E_{FRET}$ as follows:

$$E_{FRET} = \frac{1}{1 + (R/R_0)^6}$$

Here $R$ is the modeled inter-dye distance and $R_0$ is the Förster radius, which characterizes the used FRET dye pair ($R_0$ assumed constant at 54Å for the Cy3-Cy5 FRET-pair)[54]. Simulation and analysis code for the protein fingerprints are freely available at https://github.com/cvdelannoy/FRET_X_fingerprinting_simulation, while simulation data is available at https://git.wur.nl/lanno001/fret_x_proteoform_sim_data.

**In-gel proteolytic digestion using Glu-C.**

For proteomic analysis of the *O*-GlcNAc modified alpha-synuclein proteins, we performed a conventional SDS-PAGE followed by in gel proteolytic digestion and mass spectrometric analysis as previously described[45]. The aSyn proteins were analysed using a 4-12% NuPAGE Bis-Tris (Invitrogen) gel and stained with Instant Blue protein stain (Sigma) according to the manufactures instructions. The stained gel bands were cut from the gel and destained using Coomassie destaining solution (100 mM ammonium bicarbonate buffer (ABC) in 40% acetonitrile) for 15 minutes at 300 rpm at 37 ˚C. The supernatant was removed and the gel pieces were dehydrated using acetonitrile for 10 minutes at room temperatures. The supernatant was removed and the dehydrated proteins containing gel pieces were reduced using 200 µL reducing reagent solution (10mM Dithiothreitol, DTT) for 30 minutes at 56 ˚C. Next, the supernatant was removed and the samples were cooled to room temperature and the samples were alkylated for 30 minutes at RT using 200 µL alkylation reagent (55 mM Iodoacetamide in ABC buffer). After this, the alkylation solution was removed and the samples were washed with 200 µL of Coomassie destaining solution for 5 minutes at RT on a shaker. The supernatant was removed, and the samples were dehydrated using 200 µL acetonitrile for 10 minutes. Finally, 2 µL Glu-C protease stock solution (=100 ng/µL in $H_2O$, Pierce™, MS Grade) was mixed with 98 µL 100 mM ABC and added to the dehydrated gel pieces and incubated overnight at 37 ˚C under gentle shaking (300 rpm). The next day, the supernatant of each digest was collected and 150 µL of extraction solution was added to each sample and incubated for 15 minutes at 37 ˚C. The supernatant was combined with the first fraction and 100 µL of acetonitrile was added and incubated for 15 minutes at 37 ˚C, and this extract was again combined with the earlier fractions. Finally 100 µL of 10:90 ACN:$H_2O$ were added to each sample and incubated for 15 minutes at 37 °C, and combined with the earlier factions in the new Eppendorf tube. The pooled extracts from every sample were then dried using a speed-vac concentrator at elevated temperature (50–60˚C).

**β-elimination.**
To approx. 10 µL of glycopeptide/peptide extract, 300 µL of 26% dimethylamine solution were added. The reaction was carried out at 55 °C for 6 hours under careful mixing, and subsequently stopped by removing the reagent under vacuum. The residue was dissolved in 150 µL milli-Q water and stored at −20°C prior to analysis.

**Proteomic analysis.**
The speed-vac dried peptide fractions (digested or additionally beta-eliminated) were resuspended in $H_2O$ containing 3% acetonitrile and 0.01% Trifluoroacetic acid (TFA) under careful vortexing. An aliquot corresponding to approx. 100 ng digest was analysed using an one dimensional shotgun/PRM proteomics approach. Briefly, samples were analysed using a nano-liquid-chromatography separation system consisting of an EASY nano LC 1200, equipped with an Acclaim PepMap RSLC RP C18 separation column (50 µm x 150 mm, 2 µm and 100 Å), and a QE plus Orbitrap mass spectrometer (Thermo, Germany). The flow rate was maintained at 350 nL/minutes with solvent A H2O containing 0.1% formic acid, and solvent B consisted of 80% acetonitrile in H2O and 0.1% formic acid. Either a short gradient was used, consisting of a linear increase of solvent B from 5 to 30% within 38 minutes, and finally to

60% over 15 minute, or an alternative longer gradient was, consisting of a linear increase of solvent B from 5 to 25% over 88 minutes and to 55% over additional 60 minutes. In either case, the Orbitrap was operated in data-dependent acquisition (DDA) mode acquiring spectra at 70 K resolution from 385–1250 m/z, where the top 10 signals were isolated with a window 2.0 m/z for fragmentation using a NCE of 28. Fragmentation spectra were acquired at 17 K resolution, with an AGC target of 2e5, at a max IT of 75 ms. Unassigned, singly charged, 6x and higher charge states were excluded from fragmentation. Alternatively, additional (confirmatory) PRM scans were included targeting the potentially HexNAc modified peptides (inclusion list m/z = 621.3, 519.8 705.15 and 654.35).

**Processing of mass spectrometric raw data.**

Mass spectrometric raw data were analysed using PEAKS Studio X (Bioinformatics Solutions Inc., Canada) allowing 20 ppm parent ion and 0.02 Da fragment ion mass error tolerance, considering 3 missed cleavages, Carbamidomethylation as fixed and methionine oxidation and N/Q deamidation as variable modifications. The mass spectrometric raw data were furthermore analysed using a protein sequence database covering the alpha-synuclein protein sequence (synthetic, AGJ51950.1) and the GPM crap contaminant proteins sequences (https://www.thegpm.org/crap/). Additionally, decoy fusion was used for estimating false discovery rates. Peptide spectrum matches were filtered against 1% false discovery rate (FDR) and a minimum of 2 unique peptides per protein. Relative protein abundances were correlated to protein molecular weight normalised spectral counts. *O*-HexNac modified peptides were identified by including HexNAc (+203.08 Da) modifications into the variable modification search. Serine and threonine modifications sites were determined by β-elimination, where the β-elimination products were determined by including dehydration (-18.01 Da) as variable modification search. Correct annotation of HexNAc modified peptides and beta-elimination sites was ensured by additional manual investigation of identified spectra, e.g. by confirming the presence of the HexNAc oxonium ion (20.4.0872 m/z) and confirming the presence of the respective y/b peptide fragment ions from DDA and additional PRM experiments.

**Synthesis of UDP-linked 6-azido-GlcNAc.**

6-Azido-6-deoxy-N-acetyl-glucosamine-1-phosphate disodium salt was prepared as previously reported (508 mg, 1.24 mmol, 13.4% yield over 8 steps)[46]. The monophosphate was dissolved in MeOH (28 ml) and acidified to pH 5-6 by the addition of Dowex-$H^+$ resin. Resin was removed by filtration and triethylamine (4 ml) and $H_2O$ (12 ml) were added. The mixture was stirred at room temperature for 18 h and then the solvents were evaporated to obtain the triethylammonium salt of 6-azido-6-deoxy-N-acetyl-glucosamine-1-phosphate. To this compound was added trioctylamine (2.48 mmol, 1.08 ml) and the mixture was co-evaporated with pyridine (3 x 3 mL). UMP-morpholidate (1.38 g, 1.98 mmol) was added and the mixture was co-evaporated again with pyridine (3 x3 mL). The mixture was diluted with pyridine to a total volume of 12 ml and tetrazole (347.2 mg, 4.96 mmol) was added and the resulting reaction mixture was stirred at room temperature for 3 days. The reaction mixture was concentrated *in vacuo* after no more starting material was observed by TLC analysis (ethyl acetate:MeOH:$H_2O$; 4:2:1 v/v/v, staining with 10% $H_2SO_4$ in MeOH followed by charring). The crude product was purified by flash silica column chromatography (ethyl acetate:MeOH:$H_2O$, 4:2:1 v/v/v) and fractions containing carbohydrate were identified, pooled and concentrated *in vacuo*. The resulting solid was dissolved in a minimal amount of $H_2O$ and loaded on a Bio-Gel P2 Size exclusion column and eluded with $H_2O$. Fractions containing carbohydrate were identified, pooled, and lyophilized resulting in a white crystalline solid (217 mg, 0.34 mmol, 27% overall yield). $^{1}H$ NMR (600 MHz, $D_2O$) δ 7.97 (d, $J$ = 8.0 Hz, 1H), 6.00 – 5.95 (m, 2H), 5.53-5.49 (m,1H), 4.36 (dd, $J$ = 5.3, 4.5 Hz, 2H), 4.29 (m, 1H), 4.28 – 4.22 (m, 2H), 4.09 – 4.00 (m, 2H), 3.80 (app.t, $J$ = 9.8 Hz, 1H), 3.75 (dd, $J$ = 4.0, 2.0 Hz, 1H), 3.58 (app.d, $J$ = 9.1 Hz, 2H), 2.07 (s, 3H). $^{31}P$ NMR (400 MHz, $D_2O$): d −11.55 (d, $J$ = 21.4 Hz), −13.39 (d, $J$ = 21.4 Hz) ESI TOF-MS m/z calculated for $C_{17}H_{25}N_6O_{16}P_2$ $(M-H)^-$ exact 631.0802 found 631.0306.
